# Investigation of the Relationship between Markers of Systemic Inflammatory Response and Head and Neck Tumor Characteristics

**DOI:** 10.1101/399162

**Authors:** Tristan Tham, Peter Costantino

## Abstract

**Background:** Markers of systemic inflammation have been hypothesized to reflect the underlying tumor microenvironment, and have recently been shown to be associated with advanced tumor grade, T and N stages.

**Aims/Objective:** The objective of this study was to evaluate the relationship between head and neck cancer (HNC) tumor characteristics and routine pretreatment inflammatory markers: the platelet lymphocyte ratio (PLR), the neutrophil to lymphocyte ratio (NLR), and the lymphocyte to monocyte ratio (LMR).

**Materials and Methods:** This is a retrospective cohort study. The tumor characteristics collected were tumor differentiation, T stage, N stage. The relationship between the inflammatory markers and tumor characteristics was analyzed.

**Results:** A total of 122 patients were enrolled from 2010-2016. An elevated PLR was found to be significantly associated with advanced T stage (rho=0.191, p=0.00347), and N stage (ANOVA, p=0.005). None of the other inflammatory markers (NLR, LMR) were associated with T stage or N stage. No markers were associated with tumor differentiation.

**Conclusion and significance:** We found that an PLR is significantly associated with advanced tumor and nodal stage. We were unable to find any tumor associations with the other inflammatory markers (NLR, LMR).

## 1. INTRODUCTION

Head and neck cancers (HNC) accounts for a significant proportion of cancers in the general population. ^1^ These cancers are a heterogeneous group of cancers that affect the upper aerodigestive tract and the associated subsites. The current guidelines for the management of HNCs involves a variety of treatment modalities involving combinations of surgery, radiation, and chemotherapy. ^2^Treatment modality depends on the site, stage, and various other tumor or patients factors. There has been a recent paradigm shift in treating HNCs based on specific tumor related prognostic factors. One such tumor-specific factor is the human papillomavirus status (HPV) of the tumor. ^3^ Other than HPV, there has been increasing interest in other tumor or patient related factors that could impact prognosis. ^4–6^

Recently, the combination of platelet count to neutrophil-lymphocyte ratio (COP-NLR) has been reported as a prognostic indicator in head and neck cancer^7^and other sites. ^8^The study by Nakayama reported only in patients with advanced disease (Stage 3 and Stage 4 disease). Another finding in their paper was that the modified COP-NLR was a better prognostic model than the other COP-NLR models, the 4-group COP-NLR model and the Original COP-NLR model.^7^ In this context, we designed our study to report the effect of the modified COP-NLR in HNC patients of all stages.

## 2. MATERIALS AND METHODS

### 2.1 Study Design

This retrospective cohort study included patients with HNC treated at the New York Head & Neck Institute (NYHNI) from 2009 - 2016. This study was approved by the Institutional Review Board of the Northwell Health System (IRB# 17-0280-LHH). The inclusion criteria for the study was (a) histologically confirmed HNC (excluding thyroid), (b) undergoing curative intent surgery, (c) availability of complete clinical data and disease records, and (d) treatment with curative intent. The exclusion criteria was (a) clinical evidence of pre-treatment infection or other inflammatory disease or comorbidity that would alter results of the CBC, (b) benign disease (c) biopsies or non-curative intent procedures, or palliative procedures, (d) lymphoproliferative malignancies, (e) incomplete medical records, including absence of preoperative CBC, (f) CBC that was taken more than two weeks pre-operatively, (g) neoadjuvant chemotherapy or radiation therapy. Patients were screened using pre-defined ICD-9/10 codes (Supplementary materials). All patients were treated according to plans that were consistent with contemporary treatment paradigms. ^2^

### 2.2 Variable selection

Established prognostic factors for HNC were included in the data collection process: Age, Sex, BMI, Tumor site, Tumor stage according to the 8^th^ Edition of the AJCC^3^, Adjuvant treatment type, and comorbidities (ACE-27 criteria)^9^. The pretreatment modified COP-NLR was also recorded, as described by Nakayama.^7^ The NLR was defined as the neutrophil count divided by the lymphocyte count. The COP-NLR was graded as 0 for both NLR<3 and platelet count <3×10^9^/L. The COP-NLR was graded 1 for either NLR≥3 or platelet count ≥3×10^9^/L. The COP-NLR was graded as 2 for both NLR ≥3 and platelet count ≥3×10^9^/L. The electronic healthcare record of patients were reviewed and stored in a REDCAP database. ^10^ Clinical information was retrieved from the patient’s scanned notes and relevant clinical, pathological, or laboratory reports.

### 2.4 Endpoint

Two primary endpoints of this study were overall survival (OS) and event-free survival (EFS). OS was defubed as time from date of treatment to date of last follow-up or death from any cause. Patients who did not experience the event of interest (death) were censored. We choose event-free survival (EFS) as our other endpoint. A previous meta-analysis by Michiels had shown that EFS was a more effective surrogate of OS than other secondary endpoints. ^11^ EFS was defined as date of surgery to last follow-up or ‘event’, whichever was earlier. ‘Event’ was defined as any progression, local or distant recurrence, or death from any cause. ^11^ Patients who did not experience any ‘event’ were censored.

### 2.5 Statistical analysis

We used the chi-square test and analysis of variance (ANOVA) test to evaluate the differences in the relationship between the COP-NLR and clinicopathological features. The Kaplan-Meier curve (KM) was used to estimate OS and EFS, and the log-rank test was used to find survival differences between KM curves. Additionally, we used the Cox proportional hazards model (CPH) to determine the hazard ratio (HR) of the variables listed above. Univariate and multivariate analyses were performed for both OS and EFS. In generating the multivariate CPH model, significant variables in the univariate analysis were used. HRs with 95% confidence intervals (95% CI) are presented, with two-sided p-values. The alpha level was set to 0.05, and p-values less than 0.05 were considered statistically significant. All statistical analyses were performed using MedCalc for Windows, version 15.0 (MedCalc Software, Ostend, Belgium).

## 3. RESULTS

A total of 123 patients were enrolled into this study. (Table 1) The mean age of the patients was 63.5. The primary tumor sites were Oral cavity (41), Oropharynx (25), Larynx (16), Salivary gland (12), Paranasal sinus (10), Cervical lymph node or unknown primary (8), Cutaneous (9), Nasopharynx (2). Squamous cell carcinoma was the most common histological subtype (94 patients, 76%).

There were 57 patients with a COP-NLR score of 0, 8 patients with score of 1, and 58 patients with score of 2. The clinical characteristics may be found in Table 1. Tumor AJCC stage and adjuvant treatment were found to be significantly different between the COP-NLR groups. (Table 1). With regards to OS, tumor stage (HR=3.45, 95% CI: 1.27-9.34, p=0.0151) and NLR (HR=3.64, 95%CI:1.17-11.29, p=0.0254) were found to be significantly associated with survival. Both advanced stage and NLR remained independent predictors of OS in the multivariate model. With regards to EFS, tumor stage (HR=2.52, 95% CI:1.43-4.46, p=0.0015), adjuvant chemotherapy (HR=1.95, 95%CI:1.06-3.60, p=0.0319), NLR (HR=1.2573, 95%CI:1.1331-1.3952, p<0.0001) and COP-NLR score 2 vs. 0 (HR=2.5169, 95%CI:1.34-4.73, p=0.0041) were found to be significantly associated with events (Table 2 and Table 3). In order to avoid multicollinearity, we could not include both NLR and COP-NLR into the multivariate model. Of note, the HR for COP-NLR=2 was greater than the HR for NLR. In the multivariate regression for EFS, only advanced stage (HR=2.37, 95%CI:1.30-4.33, p=0.005) and COP-NLR (HR=2.25, 95%CI:1.19-4.25, p=0.0124) were found to be significantly associated with events. Kaplan meier curves were also constructed for COP-NLR and EFS, and the log-rank test between curves showed a difference between COP-NLR score of 0 and 2 (Figure 1, p=0.0114).

## 4. DISCUSSION

Through recent advances in cancer knowledge, inflammation has been discovered to be linked to the neoplastic process. ^12^ Some of the readily obtainable markers of inflammation have been shown to be significant prognostic indicators in HNC. ^4–6^ Recently, the COP-NLR has been described as a prognostic indicator as well. ^7, 8, 13^ The COP-NLR system was originally developed to combine the inflammatory data from the NLR and information from reactive thrombocytosis in the platelet count. Building on the previous work by Nakayama, this study sought to find out if the COP-NLR was prognostic in a wide variety of HNCs in various stages. The results of this study show that COP-NLR was not associated with OS. We found that COP-NLR score of 2 was associated with poorer prognosis that COP-NLR score of 0. There was no significant difference between a score of 1 and 0 in terms of EFS. Advanced cancer stage was also a significant predictor of poorer EFS in these patients, which is expected. We also found adjuvant chemotherapy to be associated with poorer EFS. In the multivariate cox regression, only advanced stage and COP-NLR score of 2 was significantly associated with poorer EFS. Of note, there were only 8 patients who had a COP-NLR score of 1, so could be a reason as to why the COP-NLR score of 1 was not found to be associated with EFS.

A high NLR score has been shown by multiple studies to be a prognostic factor for HNC. ^6^The systemic inflammatory states seen in cancer patients are thought to arise from the interaction of the tumor microenvironment. ^14^ However, the exact mechanism behind how the NLR is a prognostic factor is not well understood. ^15^ A high NLR indicates a relative increase in neutrophils over lymphocytes. Neutrophils have been known to inhibit certain lymphocytic activities such as T cell and NK cells. ^16, 17^ Conversely, lymphocyte infiltration of tumors have been shown to have improved prognosis. ^18^ Thus, a high NLR might represent a crude marker of the poorer prognosis through these inflammatory states.^6^ The mechanism behind this relationship is likely a complex pathway and remains to be fully described. ^6,15^

A recent study by Rachidi et al described that reactive thrombocytosis was associated with poorer prognosis in HNC patients, and even the administration of anti-platelet agents was associated with superior prognosis. ^19^ Platelets are known to release specific growth factors and cytokines which might be drivers of cancer growth and inflammation, such as vascular endothelial growth factor (VEGF)^20^, tumor transforming growth factor-beta.^21^ Based on this knowledge, the COP-NLR was developed and validated in various cancers. ^7, 13, 22–24^

Our study was not able to find a prognostic effect in OS, but we found that a higher COP-NLR score was associated with poorer EFS. The modified COP-NLR score used in this study might be an easily obtainable prognostic marker. However, larger studies are needed to confirm the prognostic effect, especially among the different HNC subtypes. We also found that the NLR was significant related with OS in the multivariate model. This is interesting as perhaps the NLR might be superior to the COP-NLR in certain patient populations or with regards to certain survival outcomes.

Our study has several shortcomings which we acknowledge. Firstly, this is a single center study which was conducted as a retrospective cohort study. Thus, there might be some element of bias present in the data. Secondly, we included a diverse group of patients with HNC of various subtypes, which might have their own prognostic characteristics. Third, data on HPV and p16 status for the oropharyngeal cancer patients was also unavailable for some of the cohort, and was therefore not included in the analysis. Fourth, since this was a retrospective study, patients had a variety of treatment regimens which was not standardized.

In conclusion, our study adds data to the literature in terms of prognostic value of the COP-NLR in HNC patients of all sites and all stages, which has not been reported before. We found that a COP-NLR score of 2 was associated with poorer EFS. Further large scale and perhaps prospective studies are needed to validate the findings in this report.

## 5. DISCLOSURES/FUNDING

No funding was received for this study. The authors have no financial or personal disclosures, or conflicts of interest.

## ACKNOWLEDGEMENTS

We would like to thank the following people for assistance in the data collection process: Julian Khaymovich, Caitlin Olson, Sireesha Teegala, Michael Wotman and Josephine Coury

---

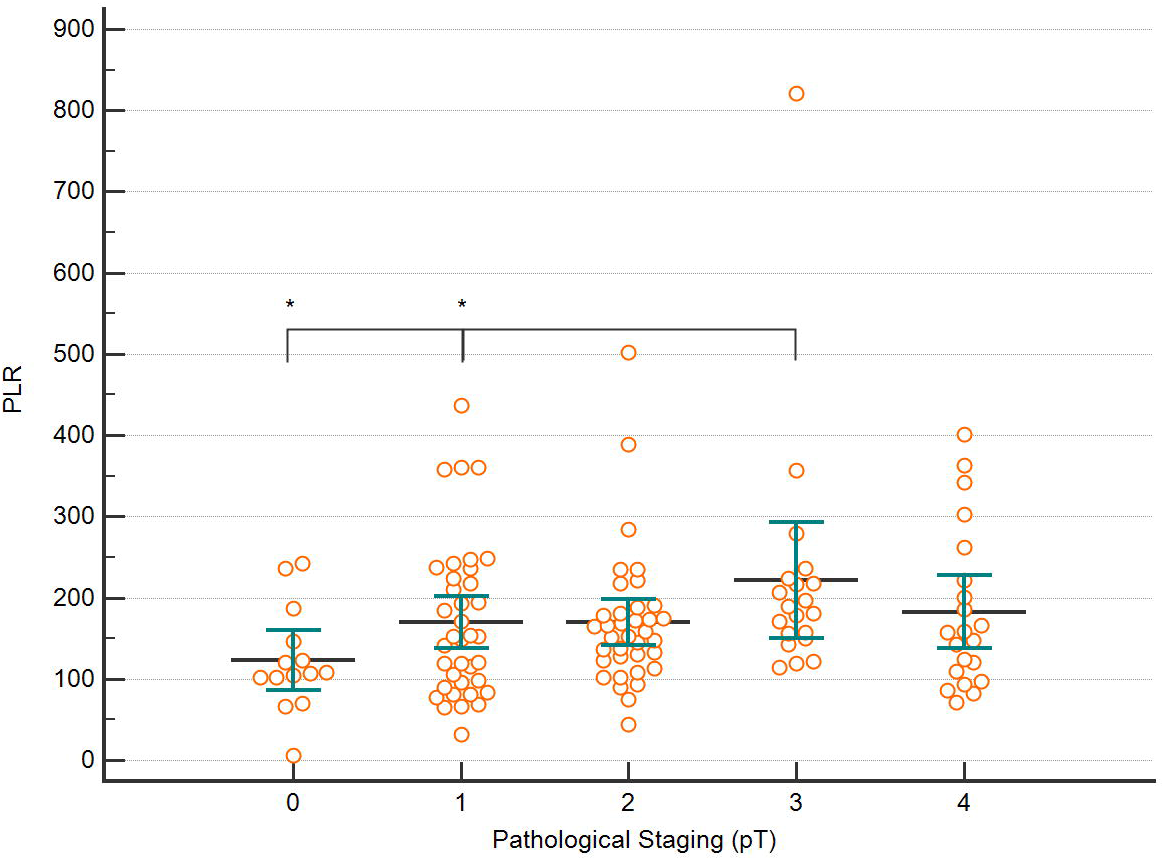

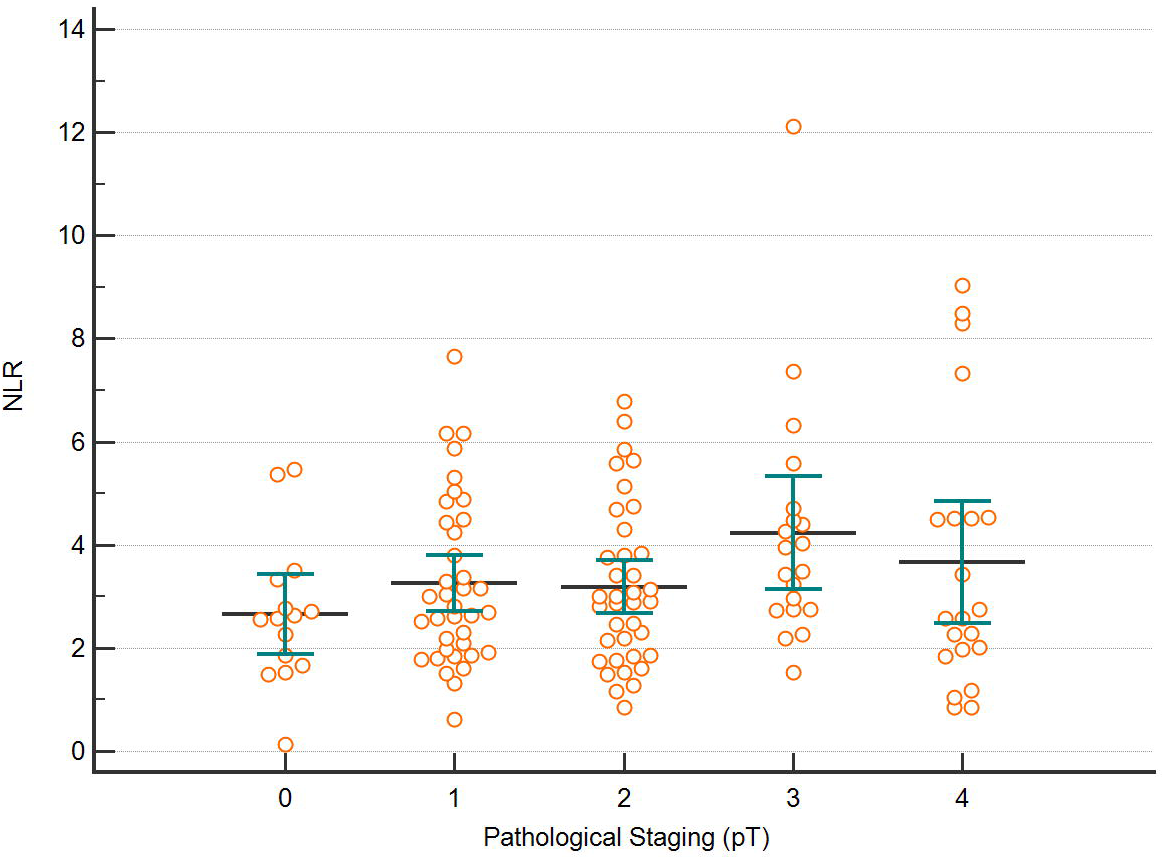

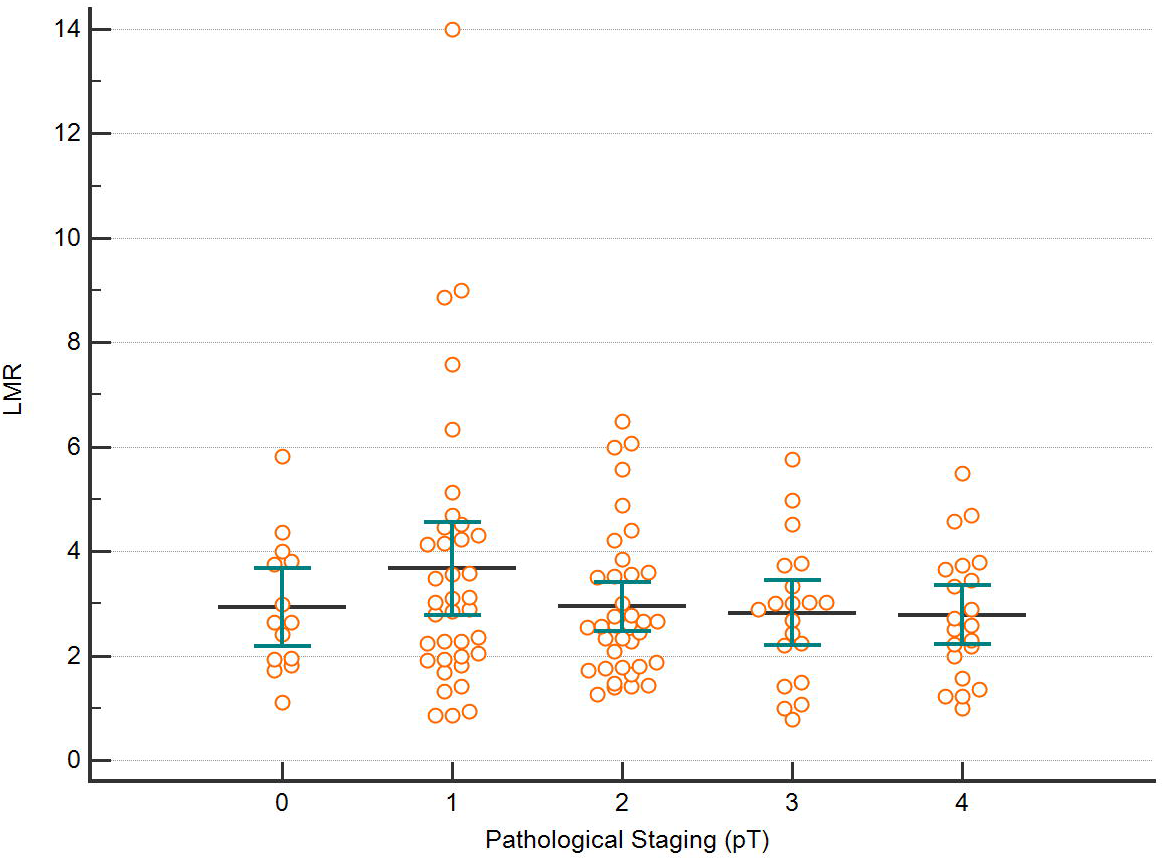

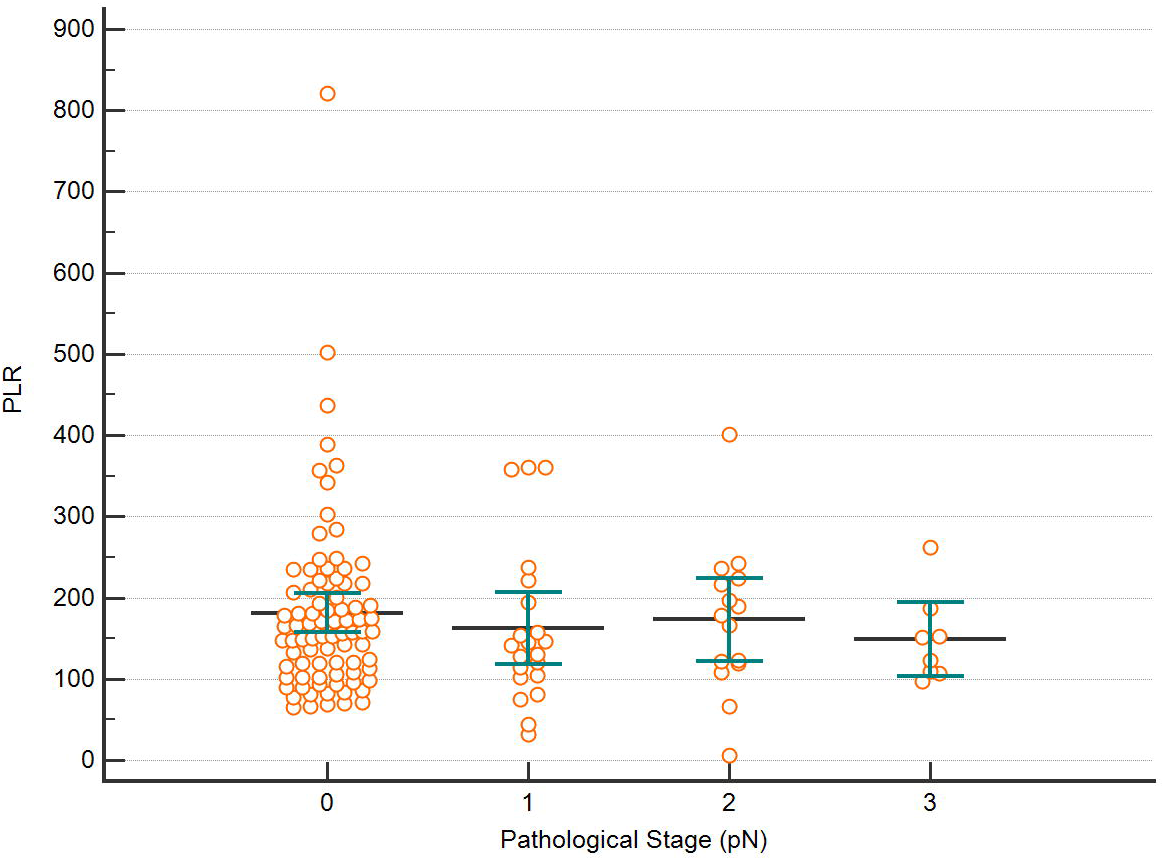

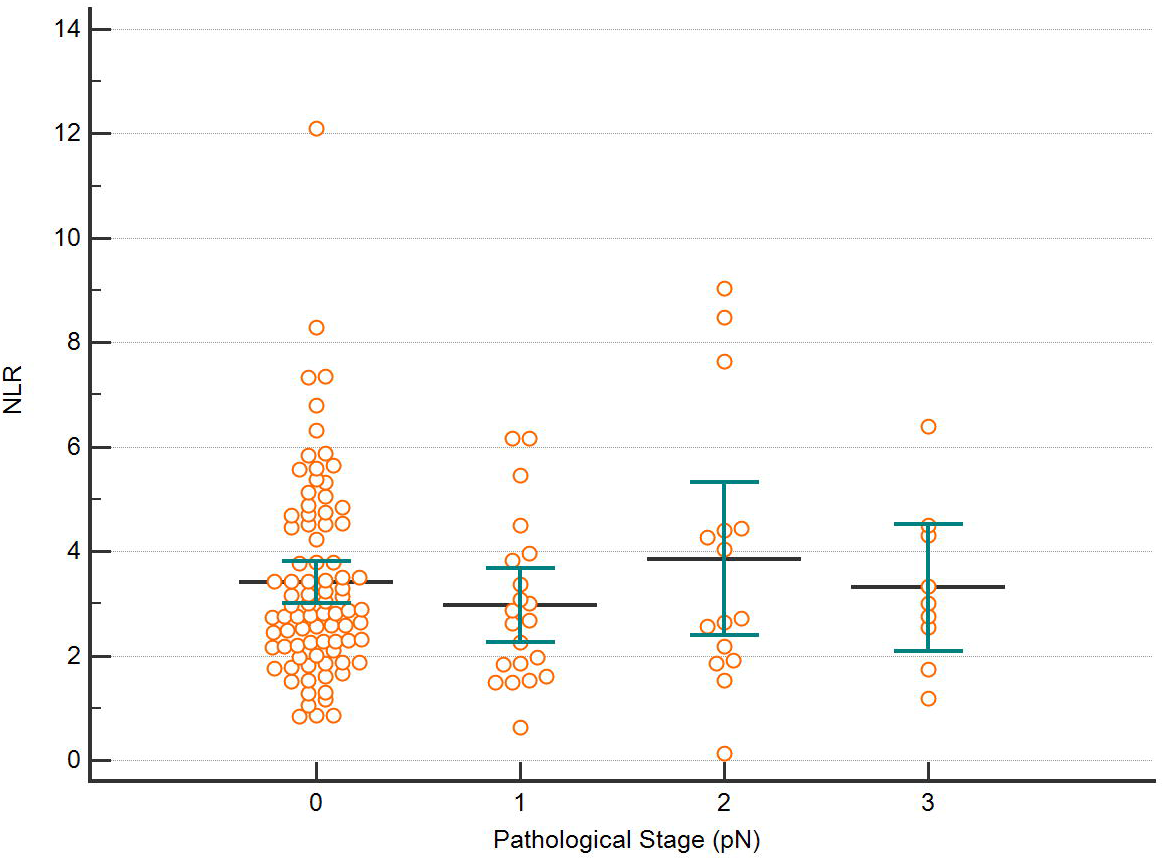

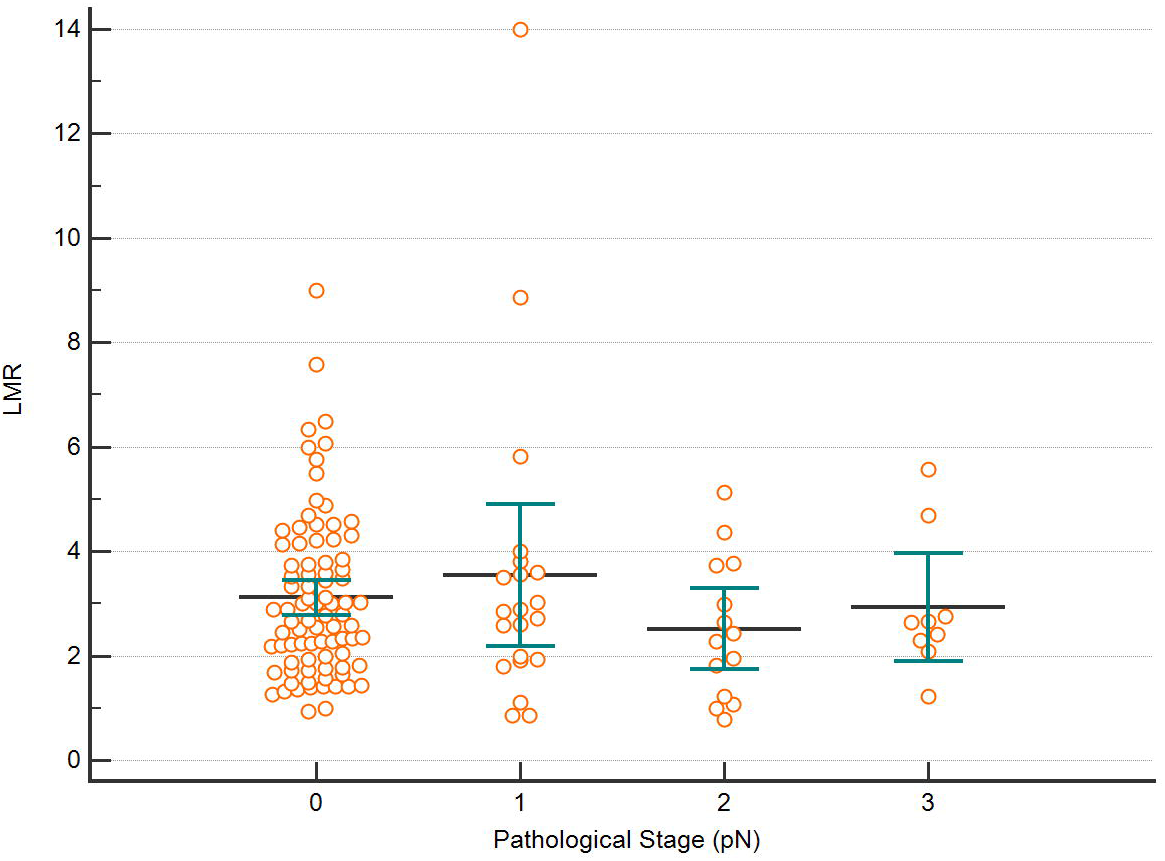

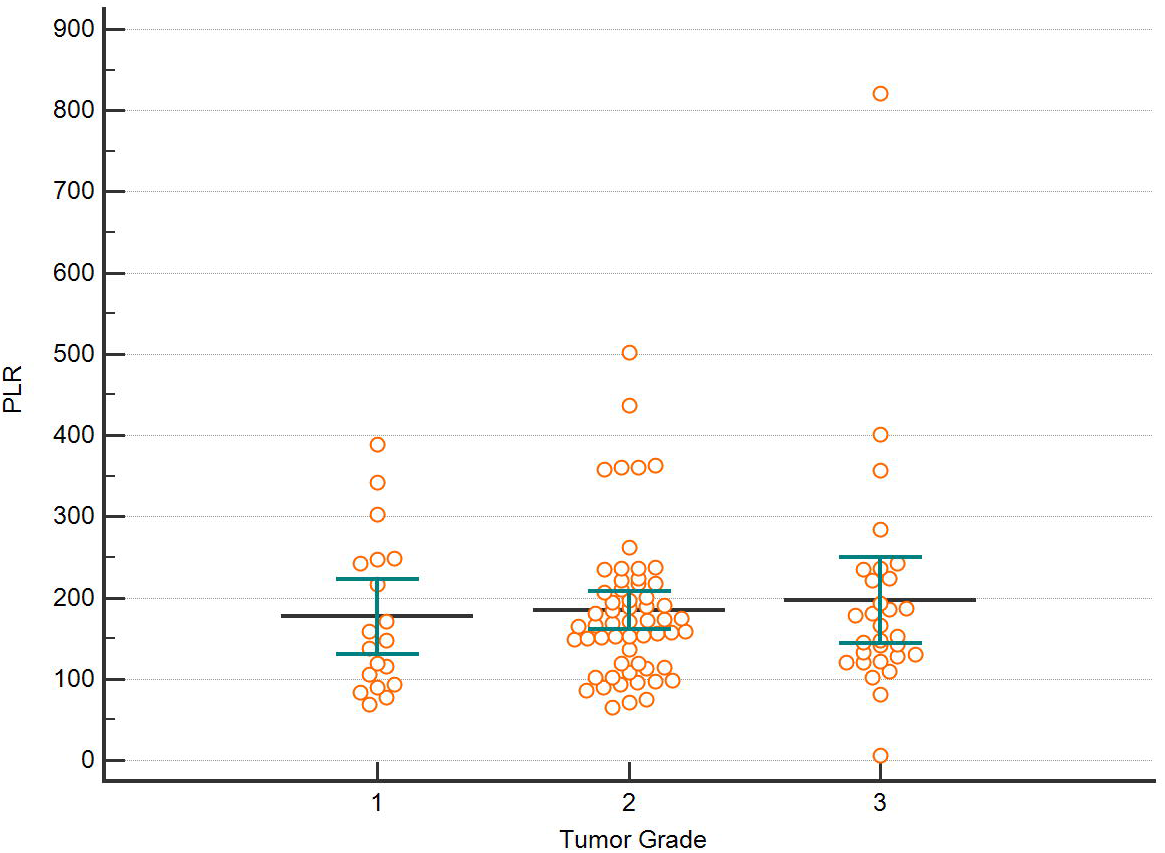

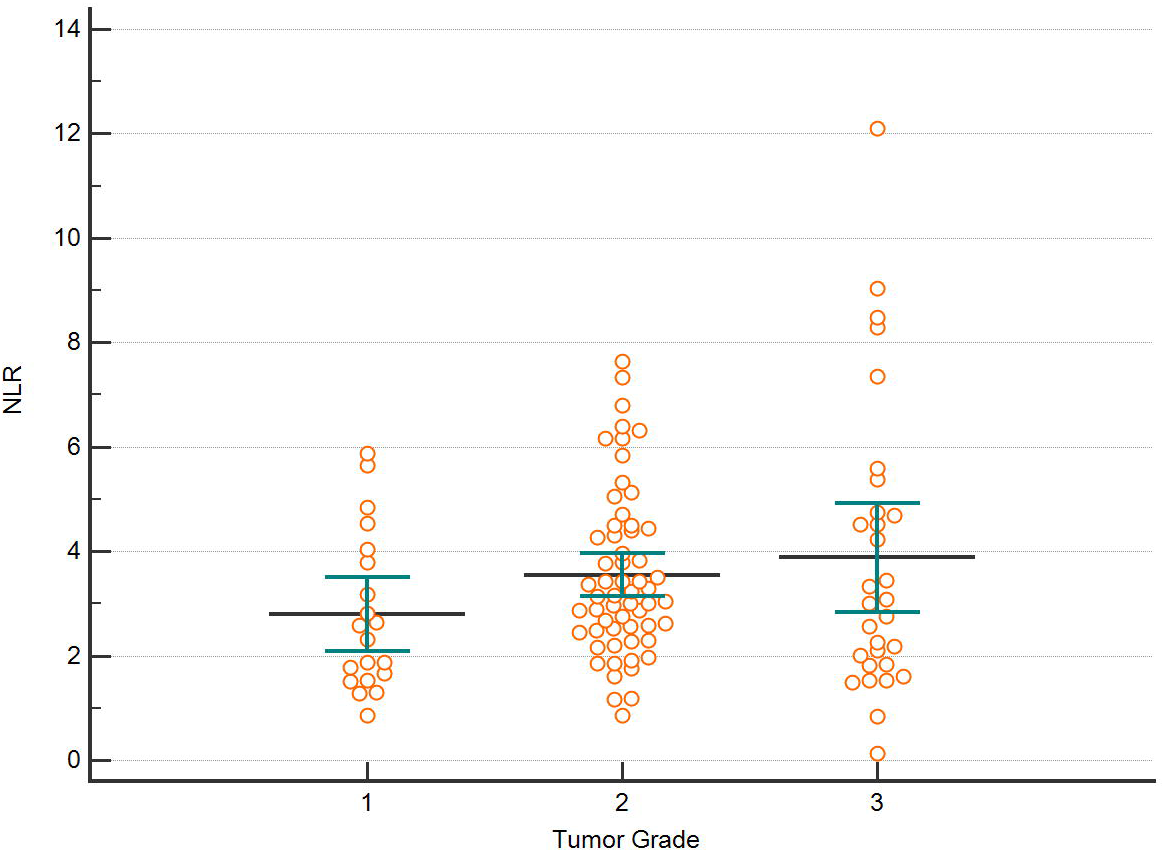

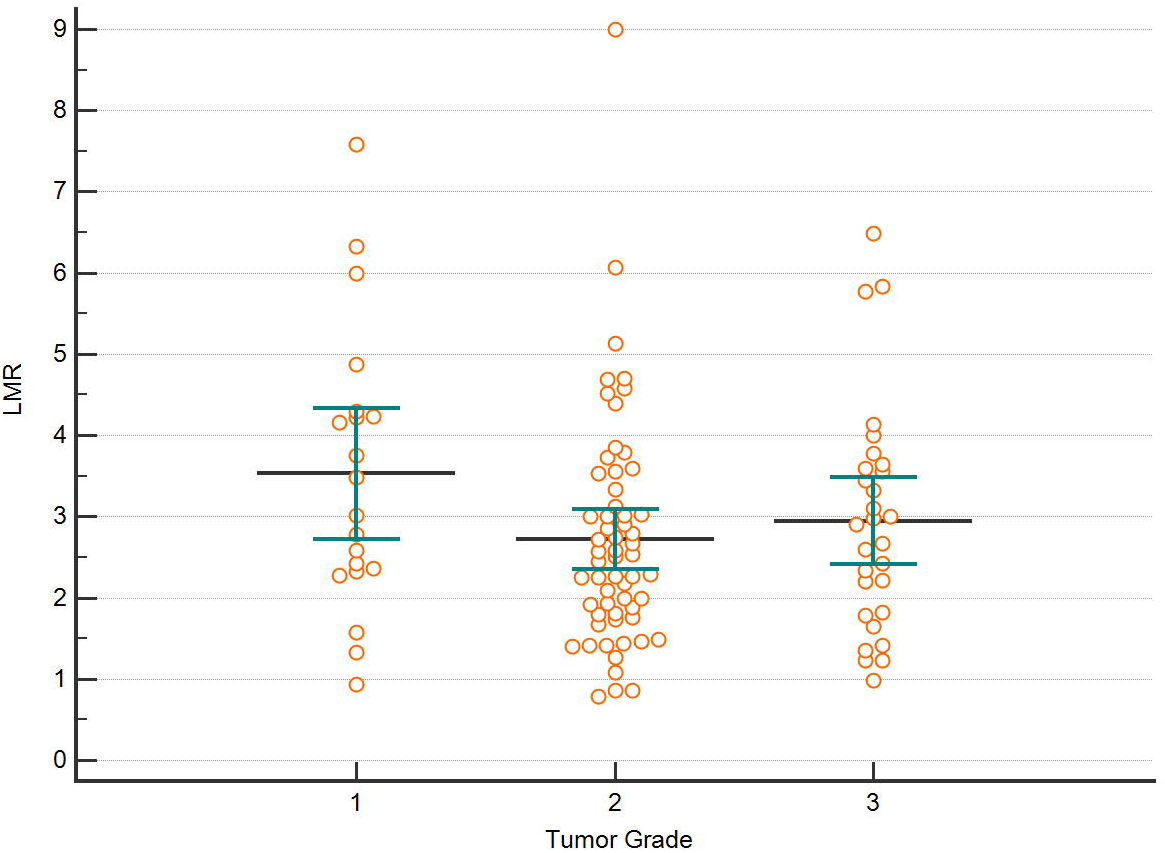

## REFERENCES

1. Jemal A, Bray F, Center MM, et al. Global cancer statistics. CA: a cancer journal for clinicians. 2011;61(2):69–90.

2. Kim HL, Puymon MR, Qin M, Guru K, Mohler JL. NCCN clinical practice guidelines in oncology™. 2014.

3. Amin MB, Edge SB, Greene FL, et al, eds. AJCC Cancer Staging Manual. 8th ed. New York: Springer; 2017.

4. Tham T, Bardash Y, Teegala S, Herman SW, Costantino PD. The red cell distribution width as a prognostic indicator in upper aerodigestive tract (UADT) cancer: A systematic review and meta-analysis. American Journal of Otolaryngology. 2018.

5. Tham T, Olson C, Khaymovich J, Herman SW, Costantino PD. The lymphocyte-to-monocyte ratio as a prognostic indicator in head and neck cancer: a systematic review and meta-analysis. European Archives of Oto-Rhino-Laryngology. 2018:1–8.

6. Tham T, Bardash Y, Herman SW, Costantino PD. Neutrophil-to-lymphocyte ratio as a prognostic indicator in head and neck cancer: A systematic review and meta-analysis. Head & Neck.0(0).

7. Nakayama M, Gosho M, Hirose Y, et al. Modified combination of platelet count and neutrophil “to” lymphocyte ratio as a prognostic factor in patients with advanced head and neck cancer. Head & neck. 2018;40(6):1138–46.

8. Tsujino T, Komura K, Ichihashi A, et al. The combination of preoperative platelet count and neutrophil lymphocyte ratio as a prognostic indicator in localized renal cell carcinoma. Oncotarget. 2017;8(66):110311.

9. Piccirillo JF, Tierney RM, Costas I, Grove L, Spitznagel Jr EL. Prognostic importance of comorbidity in a hospital-based cancer registry. Jama. 2004;291(20):2441–7.

10. Harris PA, Taylor R, Thielke R, et al. Research electronic data capture (REDCap)—a metadata-driven methodology and workflow process for providing translational research informatics support. Journal of biomedical informatics. 2009;42(2):377–81.

11. Michiels S, Le Maître A, Buyse M, et al. Surrogate endpoints for overall survival in locally advanced head and neck cancer: meta-analyses of individual patient data. The lancet oncology. 2009;10(4):341–50.

12. Grivennikov SI, Greten FR, Karin M. Immunity, inflammation, and cancer. Cell. 2010;140(6):883–99.

13. Nakahira M, Sugasawa M, Matsumura S, et al. Prognostic role of the combination of platelet count and neutrophil–lymphocyte ratio in patients with hypopharyngeal squamous cell carcinoma. European Archives of Oto-Rhino-Laryngology. 2016;273(11):3863–7.

14. Lu H, Ouyang W, Huang C. Inflammation, a key event in cancer development. Molecular cancer research. 2006;4(4):221–33.

15. Templeton AJ, McNamara MG, Šeruga B, et al. Prognostic role of neutrophil-to-lymphocyte ratio in solid tumors: a systematic review and meta-analysis. Journal of the National Cancer Institute. 2014;106(6):dju124.

16. Petrie H, Klassen L, Kay H. Inhibition of human cytotoxic T lymphocyte activity in vitro by autologous peripheral blood granulocytes. The Journal of Immunology. 1985;134(1):230–4.

17. El-Hag A, Clark R. Immunosuppression by activated human neutrophils. Dependence on the myeloperoxidase system. The Journal of Immunology. 1987;139(7):2406–13.

18. Gooden MJ, de Bock GH, Leffers N, Daemen T, Nijman HW. The prognostic influence of tumour-infiltrating lymphocytes in cancer: a systematic review with meta-analysis. British journal of cancer. 2011;105(1):93–103.

19. Rachidi S, Wallace K, Day TA, Alberg AJ, Li Z. Lower circulating platelet counts and antiplatelet therapy independently predict better outcomes in patients with head and neck squamous cell carcinoma. Journal of hematology & oncology. 2014;7(1):65.

20. Banks R, Forbes M, Kinsey S, et al. Release of the angiogenic cytokine vascular endothelial growth factor (VEGF) from platelets: significance for VEGF measurements and cancer biology. British journal of cancer. 1998;77(6):956.

21. Wakefield L, Smith D, Flanders K, Sporn M. Latent transforming growth factor-beta from human platelets. A high molecular weight complex containing precursor sequences. Journal of Biological Chemistry. 1988;263(16):7646–54.

22. Feng J-F, Huang Y, Chen Q-X. The combination of platelet count and neutrophil lymphocyte ratio is a predictive factor in patients with esophageal squamous cell carcinoma. Translational oncology. 2014;7(5):632–7.

23. Ishizuka M, Nagata H, Takagi K, Iwasaki Y, Kubota K. Combination of platelet count and neutrophil to lymphocyte ratio is a useful predictor of postoperative survival in patients with colorectal cancer. British journal of cancer. 2013;109(2):401.

24. Ishizuka M, Oyama Y, Abe A, Kubota K. Combination of platelet count and neutrophil to lymphocyte ratio is a useful predictor of postoperative survival in patients undergoing surgery for gastric cancer. Journal of surgical oncology. 2014;110(8):935–41.

